# Morph-dependent nematode infection and its association with host movement in the land snail *Cepaea nemoralis* (Mollusca, Gastropoda)

**DOI:** 10.1101/2022.03.04.482990

**Authors:** Maxime Dahirel, Marine Proux, Claudia Gérard, Armelle Ansart

**Affiliations:** Université Côte d’Azur, INRAE, CNRS, ISA, F-06903, Sophia-Antipolis, France; Department of Biology, Ghent University, B-9000, Ghent, Belgium; Univ Rennes, UR1, CNRS, ECOBIO (Ecosystèmes, Biodiversité, évolution), UMR 6553, F-35000, Rennes, France

**Keywords:** activity, *Brachylaima*, food intake, *Riccardoella*, shell colour polymorphism, snail host-parasite interactions

## Abstract

Host behaviour can be influenced by parasitic risk and infection through a variety of direct and indirect mechanisms. We can expect individuals expressing different phenotypes to also differ in the ways their behaviour is altered by parasites. We used the land snail *Cepaea nemoralis*, a species with various shell colour morphs differing in behaviour and physiology, as a model to study the link between parasite response and individual behaviour variation. We analysed metazoan parasite abundance and its relation to behaviour (movement and food intake) in snails from three shell morphs (from light unbanded to darker five-banded) and from two neighbouring populations from contrasted environments. Snails were parasitized by mites, trematodes and nematodes, from rarest to most frequent. We confirm that terrestrial gastropods can defend against infection by trapping parasitic nematodes in their shell. We show that nematode encapsulated in shells can uncover past infections even when a snail population is currently nematode-free, possibly due to parasite seasonality. We present new observations suggesting that not only nematodes but also mites might be encapsulated in shells. Infection levels varied between morphs, with darker snails harbouring fewer nematodes. Behaviour (movement and food intake) was linked to nematode, but not trematode infection. Individuals with higher nematode load ate less, irrespective of morph and population. The most-infected morph (unbanded snails) showed reduced activity in the nematode-infected population compared to the one that was nematode-free at sampling time. As a result, parasites may cancel previously described behavioural differences between morphs. We discuss the possible mechanisms explaining morph-dependent responses to parasites, and how parasite risk may be an unseen force shaping *C. nemoralis* morph variation in space and time. We conclude on the possible ecological consequences of the link, mediated by shell colour, between thermal and immune responses.

## Introduction

Parasites and the behaviour of their hosts are interlinked in complex ways: parasites change the environmental context in which potential hosts express their behaviour, and directly alter actual hosts’ internal phenotypic state; host behaviour may favour or hinder parasite spread. Focusing on movement-related behaviours (Binning, Shaw & Roche, 2017 for a review), potential hosts may react to the perceived risk of infection by moving to avoid contact with parasites (e.g. Davies & Knowles, 2001). Infection costs may force infected individuals to limit or delay movements (e.g. Debeffe *et al*., 2014; O’Dwyer, Kamiya & Poulin, 2014), possibly reshaping the distribution of infection risk in space. All these responses may be shaped by pre-existing behavioural variation, leading to feedbacks between parasite dynamics and the behaviour of uninfected and infected individuals (Ezenwa *et al*., 2016). To untangle these dynamics, and understand their consequences, we need to map the existing relationships between parasite infection and behavioural variation.

Land snails and slugs host a diverse array of metazoan parasites and parasitoids, including flies, mites, cestodes, trematodes and nematodes (Barker, 2004; Morley & Lewis, 2008; Gracenea & Gállego, 2017; Pieterse, Malan & Ross, 2017; Gérard *et al*., 2020). Depending on species, parasites use terrestrial gastropods as their only hosts, or as intermediate hosts with definitive hosts often being birds and mammals (Barker, 2004). Some of these parasites can cause diseases (e.g., strongylosis) in domesticated animals and humans (Meerburg, Singleton & Kijlstra, 2009; Butcher, 2016; Giannelli *et al*., 2016). Parasite infection is often highly variable between populations (Baur & Baur, 2005; Żbikowska *et al*., 2020).

Terrestrial gastropods can defend against metazoan parasites in several ways. They may alter their behaviour to avoid parasite-infected microhabitats (Wynne, Morris & Rae, 2016). If parasites manage to enter snails, they can be neutralized through several physiological mechanisms, including cellular and non-cellular immune responses (Furuta & Yamaguchi, 2001; Coaglio *et al*., 2018). Snails can also trap nematodes directly in the inner layer of their shell (Rae, 2017). As in insects (González-Santoyo & Córdoba-Aguilar, 2012; San-Jose & Roulin, 2018), melanin and the phenoloxidase pathway involved in melanisation play a key role in snail defences against parasites (Coaglio *et al*., 2018). This association between immunity and pigment-related pathways could be especially relevant in land snails. Indeed, many species exhibit shell colour polymorphism (Williams, 2017), which affects behaviour, thermal responses and vulnerability to predators (e.g. Jones, Leith & Rawlings, 1977; Rosin *et al*., 2011; Surmacki, Ożarowska-Nowicka & Rosin, 2013; Köhler *et al*., 2021).

The grove snail *Cepaea nemoralis* has a long history as an iconic model in evolutionary biology (Jones *et al*., 1977; Ożgo, 2009). Individuals can differ in background shell color (yellow, pink or brown) as well as in the number of dark bands (zero to five). Decades of research have aimed to disentangle the roles of thermal selection, predation risk and neutral processes in driving the widely observed, across multiple scales, among-population differences in morph frequencies (Cain & Sheppard, 1954; Lamotte, 1959; Jones *et al*., 1977; Schilthuizen, 2013; Ożgo *et al*., 2017). These color differences are associated with behavioural and physiological differences (e.g. Kavaliers, 1992; Dahirel, Gaudu & Ansart, 2021), leading to potentially adaptive phenotypic syndromes.

If susceptibility and response to parasite infection are shaped by host behaviour and physiology, this means that (i) *C. nemoralis* morphs might consistently differ in their infection levels, (ii) variation in infection between populations may alter host behaviour in a morph-dependent way, (iii) infection and behaviour may also covary independently of morph. In this context, we compared natural infections by metazoan parasites across two habitats differing in microclimate conditions. We examined whether infections varied with morph and potentially morph-dependent behaviours, namely food intake and activity.

Based on the link between pigments and immune defence outlined above, we expected snails with more dark bands hosted fewer parasites. Since darker snails are less mobile (Dahirel *et al*., 2021), this is in line with some predictions from the pace-of-life syndrome hypothesis, namely that fast active individuals have lower immune responses (Réale *et al*., 2010; but see Royauté *et al*., 2018). We also hypothesized that metazoan parasites would have notable negative effects on snail performance (McElroy & de Buron, 2014), leading to slower behaviours and changed food intake in more infected snails (both at the morph level and within morphs). We expected these effects on behaviour to be stronger in morph(s) more vulnerable to parasite infection.

## Methods

### Snail collection and maintenance

We collected snails in early May 2019 (i.e. at most ≈ 1.5 month after hibernation) from two contrasted sites in and near the village of Arçais, France. From previous studies, we know that snail parasites are present, albeit in another species (*Cornu aspersum*, Gérard *et al*., 2020), and that environmental differences between the sites are linked to differences in the frequency of darker *Cepaea nemoralis* morphs (Dahirel *et al*., 2021).

- The “open habitat” population is in a garden with little to no trees and thus exposed to the sun throughout the year (≈ 46°17’50”N, 0°41’30” W).
- The “shaded habitat” population (≈ 46°18’01” N, 0°42’56” W) is in a small humid woodlot which is not directly exposed to the sun during snail activity months, and occasionally experiences partial winter flooding. The substrate is poorer in calcite than in the open habitat (Dupuis *et al*., 1983; BRGM, 2005). Five-banded snails are more frequent there, in agreement with hypotheses on shell colour and thermal selection (22.3% of individuals, compared to 13.5% in the open habitat; Dahirel *et al*., 2021).

We sampled 90 adult individuals per population, divided equally into unbanded, three-banded, and five-banded snails. Adults were recognizable by the presence of a reflected “lip” around the shell opening. We only sampled snails with yellow background shells, leaving aside the rarer brown and pink-shelled morphs. Sampling was random with respect to other shell features (e.g. band fusion).

We kept snails under standard conditions (20 ± 1°C, L:D 16:8) in 30 × 45 × 8 cm^3^ plastic crates lined with 1-2 cm of soil kept humid and regularly renewed, with a lid made of a fine plastic mesh. Snails were grouped by morph × population of origin. Food (vegetal flour supplemented with calcium, Hélinove, Saint Paul en Pareds, France) was available *ad libitum* in Petri dishes.

Four days before the experiments, we marked snails with a unique identifier on the side of their shell (using Posca pens, uni Mitsubishi Pencil, Tokyo, Japan). At the same time, we measured snail size (greater shell diameter) to the nearest 0.1 mm.

### Movement behaviour and food intake

Behavioural experiments began two weeks after snail sampling. We tested each snail twice for both movement and food intake, to separate repeatable among-individual variation from within-individual variability. To ensure a manageable workload, we split the 180 snails in three groups of 60, each starting the experimental sequence on a different day, with each group containing snails from all band number × population combinations in equal proportions.

We first tested snails for movement behaviour. We stimulated snail activity by placing them in a Petri dish containing ≈ 1 mm of water until they were out of their shell. We then placed each snail individually on a clean lab bench, and used the time snails took to move more than 10 cm in a straight line from their starting point as a measure of movement (longer times implying slower individuals). Importantly, we only counted time after snails moved more than 2 cm (i.e. about one shell length) from their starting point, to ensure all observed snails were truly active. Some snails stayed inactive and the relevant observations were thus excluded (11 individuals missed one of the two tests, 6 individuals missed both). We stopped observation 20 min after placing snails on the lab bench (not after the start of activity).

Movement tests were immediately followed by food intake tests. We kept all snails individually in 10 × 7 × 6 cm^3^ boxes for 18 hours (including 8 hours of darkness) and provided them with 1.5 g of dry food (a non-limiting quantity, large enough to feed one snail for at least three days, based on preliminary tests). Afterwards, we recovered, dried (60°C for 60 hours) and weighed the remaining food (to the nearest 0.1 mg) to estimate food intake.

Snails were then placed back in their original box. The second test for each behaviour was done one week later.

### Parasitological research

After freezing at −20°C and thawing, we dissected each snail under a binocular microscope. After removing the body from the shell, we looked for and counted metazoan parasites in various locations: space between shell and soft body, lung, heart, kidney, body cavity and digestive gland, as in Gérard et al. (2020). We found nematodes, larval trematodes and parasitic mites (Acari). Trematodes were identified to the genus level as in Gérard et al. (2020), and mites using Fain (2004). That level of taxonomic precision could not be reached for nematodes. However, nematodes from different families consistently use/infect different organs and reach different body sizes within snails (Morand, Wilson & Glen, 2004; Wilson, 2012). We assigned nematodes to size classes (<1 mm, 1-3 mm, 3-5 mm, >5mm) using graph paper, and used nematode location and size to infer the most likely families infecting our snails. We additionally observed shells and shell fragments under brighter light to detect encapsulated parasites (Rae, 2017). By simplicity, we use “live” as a shorthand to designate parasites found free within the body or between the body and the shell during dissection, and thus presumably still living before snails were sacrificed, by opposition with “trapped” or “encapsulated” parasites found within shells.

### Statistical analyses

We analysed our data in a Bayesian framework, using R (version 4.2.0, R Core Team, 2022) and the brms R package (Bürkner, 2017) as frontends for the Stan language (Carpenter *et al*., 2017). We used the tidybayes, bayesplot, patchwork packages for data preparation, model evaluation and plotting, as well as the tidyverse suite of packages (Gabry *et al*., 2019; Kay, 2019; Wickham *et al*., 2019; Pedersen, 2020). One snail (out of 180) died during the experiments and was excluded from statistical analyses, but was used for natural history information (see **Results**).

We ran two models, which both included at least effects of population of origin, number of shell bands, as well as population × banding interactions as categorical explanatory variables (hereafter “morph and population effects”).

We first used a Gaussian linear model to analyse how snail shell diameter (mean centred and scaled to unit 1 SD), varied with morph and population effects.

We then ran a multivariate generalized mixed/multilevel model to examine how behaviour and parasite infections varied between morph and populations, and whether these traits were correlated at the individual level. Trematodes and nematodes could not be identified to species, so we used the total numbers of trematodes and nematodes per host snail in our analyses (as in e.g. Debeffe *et al*., 2014). Mite infections were too rare to be analysed statistically **(Results).** This multivariate model included four submodels:

- live trematode abundance was modelled as a function of morph and population effects, as well as shell size, using a Poisson model. This model included random effects of test group and individual identity. The latter acts as an observation-level random effect and accounts for overdispersion (Harrison, 2014); it also allows us to build an individual-level correlation matrix connecting all four traits.
- live nematode abundance was modeled as trematode abundance, with one change. As no live nematodes were found in the shaded population, we only fit this submodel to data from the open population and removed the population fixed effect. Importantly, keeping all data instead does not change our conclusions (see **Data availability**).
- food consumption (mean-centred and scaled to unit 1SD) was analysed using a Gaussian model including the same fixed and random effects as trematode abundance. Although food consumption data are proportions (of total food), and so are well-suited to Beta models (Douma & Weedon, 2019), the Gaussian model behaved much better regarding posterior predictive checks **(Data availability).** As individuals were measured twice, the individual random effect here allows separation between within- and among individual variation.
- movement behaviour (latency to move more than 10 cm from the starting point) was inverse-transformed, mean-centred and scaled to unit 1SD, and then analysed using a Gaussian model including the same fixed and random effects as above. We initially attempted to analyse untransformed latencies using lognormal or Gamma models, but the Gaussian model performed better regarding posterior predictive checks (**Data availability**).

In both models we used weakly informative priors (McElreath, 2020): Normal(0,1) priors for fixed effects, half – Normal(0,1) priors for standard deviations (random effect and residual), and a LKJ(2) prior for the correlation matrix in the multivariate model. See **Supporting Information S1** for detailed model descriptions. We ran four chains per model for 4000 iterations each, with the first half of each chain used as warmup. We checked convergence and mixing following Vehtari et *al*. (2021). All parameters had satisfactory effective sample sizes (Vehtari *et al*., 2021). Posteriors are summarised in text and figures as means [95% Highest Posterior Density Intervals].

We also looked at the proportion of snails with encapsulated nematodes as a function of shell morph and population. Here we did not run statistical models, and simply present these proportions as qualitative information about past parasite exposure in both populations. We explicitly refrained from analysing the prevalence or abundance of encapsulated nematodes at the individual level within populations, as it is likely linked to cumulative exposure time/age, and snail age is not known (and cannot be approximated by e.g. size, given *Cepaea nemoralis* exhibits determinate growth).

## Results

### Snail size

Snails from the shaded population were on average smaller than those from the open habitat if they were three-banded (*Δ*_[shaded–open]_ = −0.70 mm [−1.38; −0.04], Fig. 1) or five-banded (*Δ*_[shaded–open]_ = −1.37mm [−2.07; −0.71], Fig. 1). For unbanded snails, there was no clear difference in size between habitats (*Δ*_[shaded–open]_ = −0.39 mm [−1.03; 0.33], Fig. 1).

**Figure 1.**
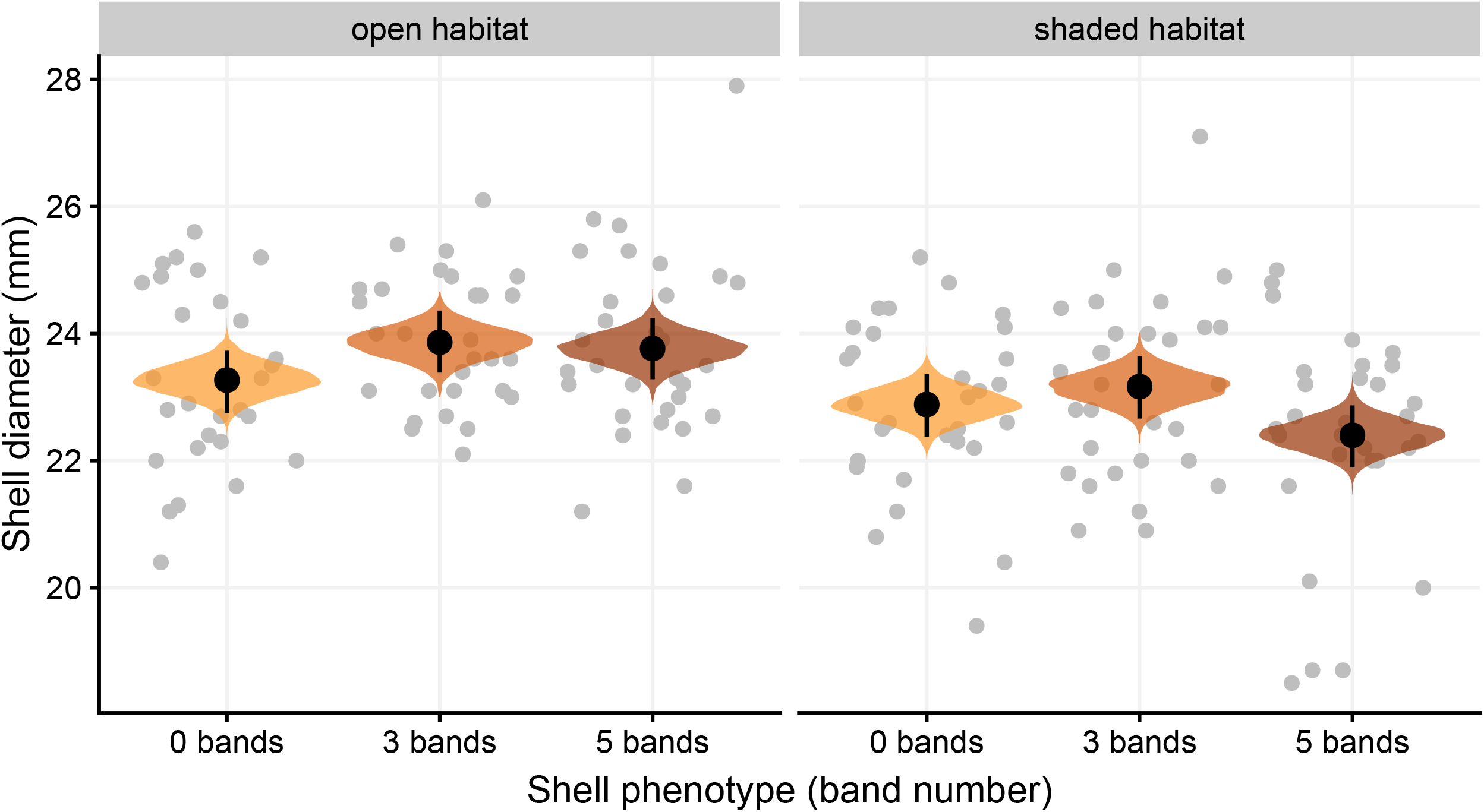
Snail shell diameter as a function of band number and population of origin. Grey dots are observed values, displayed along the posterior distributions of the average sizes (black dots and segments are the means and 95% CI of the posteriors).

### Parasitological information

Snails from both populations and all morphs were infected by metazoan parasites **(Figs. 2-3,** see **Supporting Information S2** for detailed descriptive statistics about prevalence and infection intensity). All trematodes found were live *Brachylaima* sp. metacercariae in snails’ kidneys; no live sporocysts were recorded and no trematodes of any stage were found trapped in shells. Among the organs we examined, we detected live nematodes only within the lung and in the space between the body and the shell; most of these nematodes were in the 1-3 mm size class (297/371; 211/269 excluding the snail that died during the experiments). Based on this, we can tentatively assume that most nematodes found belonged to the Rhabditidae family (Morand *et al*., 2004; Wilson, 2012). In addition we frequently (Fig. 3) found nematodes encapsulated in shells. No live nematodes were found in individuals from the shaded population **(Fig.3),** by contrast to the open population where 65.6% of snails were infected (59/90, including the dead snail). However, a high number of snails contained encapsulated nematodes in their shells, in both populations **(Figs 2, 3),** indicating past exposure to nematodes even in the shaded population. Only one snail (unbanded from the shaded habitat) harboured live mites (*N* = 93 mites), identified as *Riccardoella* sp. Three other snails (all five-banded from the shaded habitat) had (unidentified) mites trapped in their shell (*N* = 1, 2 and 15 mites, **Fig. 2).** As mentioned in **Methods,** one snail among 180 died during the experiments; it harboured 102 live nematodes, more than 4 times the number of the most infected surviving snail **(Fig. 3)**.

**Figure 2.**
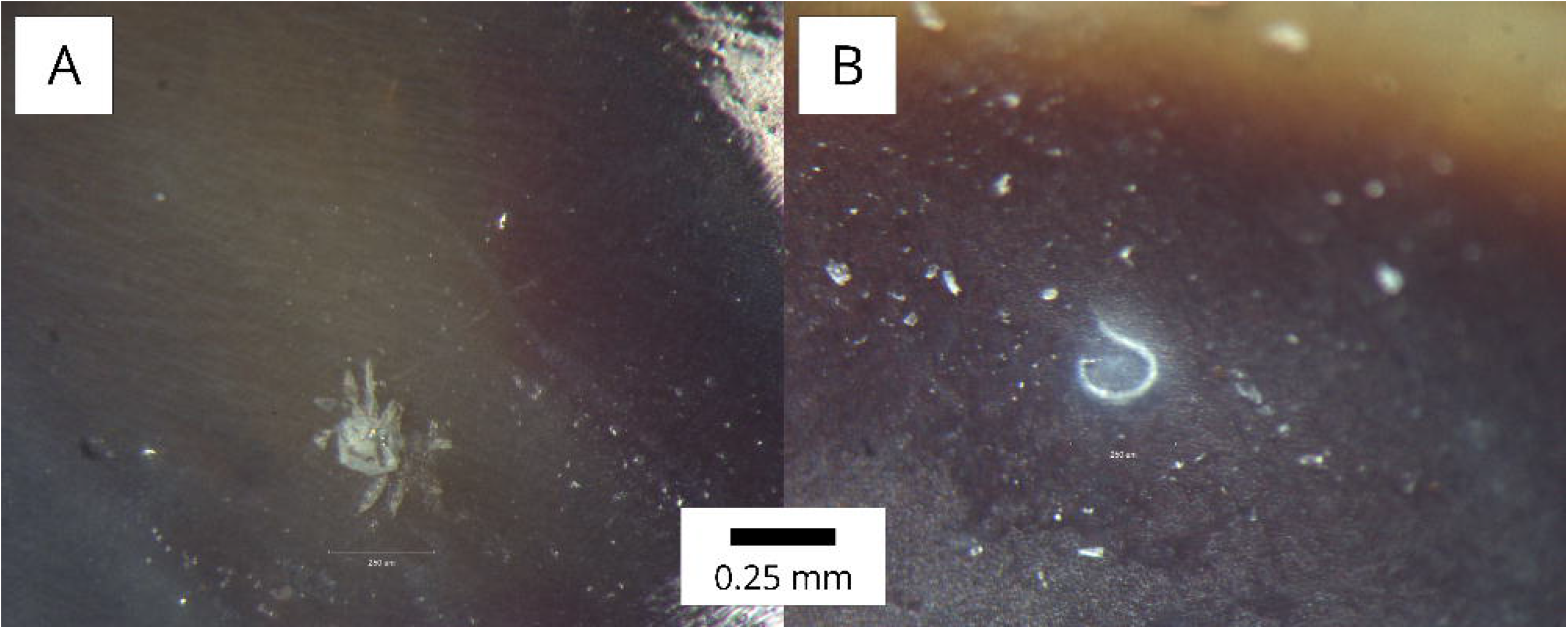
Examples of encapsulated parasites in *Cepaea nemoralis* shells. (A) mite; (B) nematode. Original images, including other images of encapsulated parasites, are made available with the rest of the data (see **Data availability**).

**Figure 3.**
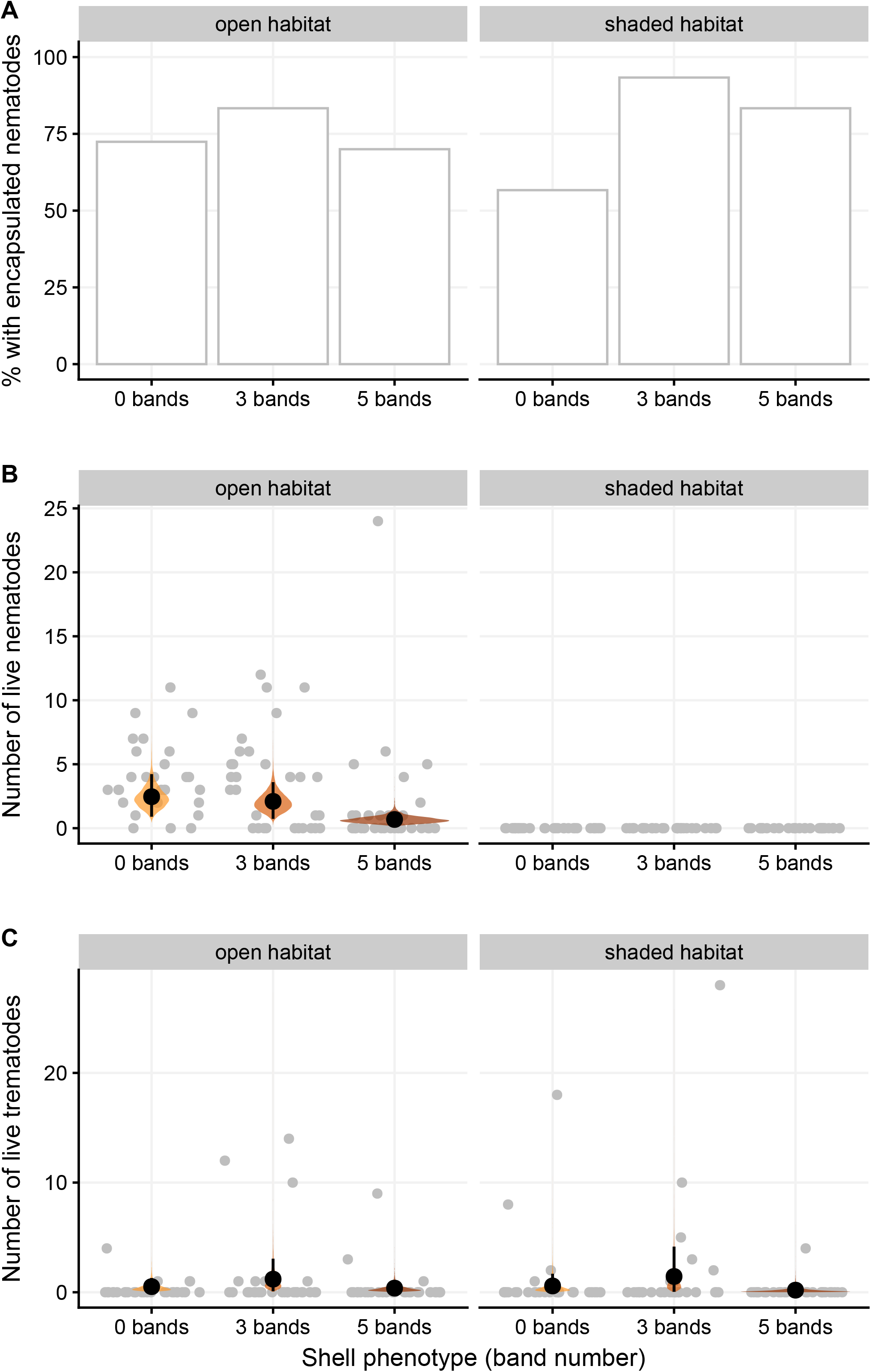
Parasite-related variables as a function of morph and population of origin (*N* = 179, one dead snail was excluded). (A) Proportion of snails having at least one nematode trapped in their shells; (B) Live nematode abundance; (C) Live *Brachylaima* sp. trematode abundance (metacercariae). For (B) and (C): grey dots are observed values, displayed along the posterior distributions of the average abundances based on the model described **Table 1** (black dots and segments are the means and 95% CI of the posteriors). Posterior predictions are based on fixed effects only, and the effects of shell size are averaged out. See **Supporting Information S3** for pairwise comparisons.

In the open population where live nematodes were found, nematode abundance was morph-dependent: five-banded snails were on average infected by about a third of the nematodes infecting other snails (*N*_[5bands]_/*N*_[0bands]_ = 0.29 [0.11; 0.52], *N*_[5bands]_/*N*_[3bands]_ = 0.34 [0.12; 0.62]; **Table 1, Fig. 3),** mostly due to differences in prevalence between morphs **(Supporting Information S2).** Five-banded snails also contained fewer trematodes, but only when compared to three-banded snails and only clearly in the shaded habitat (*N*_[5bands]_/*N*_[3bands]_ = 0.19 [0.00; 0.57]; **Fig. 3, Supporting Information S3).** Parasite abundance (trematode or nematode) was not linked to shell size **(Table 1**).

**Table 1.** Summary of posterior model coefficients for the parasite and behaviour multivariate model. Individual-level correlations between traits are presented in **Table 2.** See **Methods** for model details, and **Supporting Information S3** for pairwise comparisons among morph × population groups. Fixed effects of population and size that are different from 0, based on 95% intervals are in bold.

|  | Nematode<br>abundance* | Trematode<br>abundance | Movement | Food intake |
| --- | --- | --- | --- | --- |
| **Fixed effects** |  |  |  |  |
| Intercept<sub>[0 bands]</sub> | 0.84 [0.09; 1.52] | -0.94 [-2.44; 0.48] | -0.26 [-0.66; 0.16] | -0.22 [-0.61; 0.19] |
| Intercept<sub>[3 bands]</sub> | 0.67 [-0.08; 1.38] | -0.10 [-1.54; 1.33] | 0.16 [-0.24; 0.56] | -0.09 [-0.50; 0.31] |
| Intercept<sub>[5 bands]</sub> | -0.47 [-1.31; 0.32] | -1.26 [-2.72; 0.23] | -0.09 [-0.50; 0.31] | -0.04 [-0.45; 0.37] |
| Population = “shaded”<sub>[0 bands]</sub> | — | -0.02 [-1.42; 1.39] | **0.71 [0.31; 1.08]** | **0.42 [0.04; 0.83]** |
| Population = “shaded”<sub>[3 bands]</sub> | — | 0.06 [-1.31; 1.34] | -0.23 [-0.61; 0.13] | 0.22 [-0.18; 0.63] |
| Population = “shaded”<sub>[5 bands]</sub> | — | -0.88 [-2.44; 0.66] | -0.08 [-0.47; 0.32] | 0.13 [-0.26; 0.55] |
| Shell size (standardised) | -0.19 [-0.53; 0.13] | 0.06 [-0.57; 0.72] | 0.09 [-0.03; 0.21] | **0.23 [0.10; 0.35]** |
| **Random effects SDs** |  |  |  |  |
| Test group SD | 0.53 [0.00; 1.23] | 1.95 [0.94; 3.19] | 0.23 [0.00; 0.70] | 0.21 [0.00; 0.67] |
| Individual-level SD | 1.14 [0.85; 1.46] | 2.57 [1.99; 3.21] | 0.32 [0.04; 0.54] | 0.56 [0.43; 0.70] |
| **Residual SD** | — | — | 0.92 [0.82; 1.02] | 0.8 [0.72; 0.88] |

\* The nematode model is only estimated for the “sun-exposed” population, since no live nematodes were recorded in the “shaded” population

### Snail behaviour

Both movement and food intake were repeatable at the individual level (see individual-level SDs **Table 1**), although adjusted repeatability (sensu Nakagawa & Schielzeth, 2010) was weaker and closer to 0 for movement (*r*_[movement]_ = 0.11 [0.00; 0.23]; *r*_[food intake]_ −0.29 [0.15; 0.42]; **Supporting Information S4)**.

Movement depended on both population of origin and shell band number **(Table 1, Fig. 4, Supporting Information S3).** In the shaded population, five-banded snails were on average less mobile than unbanded snails (*Δ*_[5bands–0bands]_ = −0.61 SDs [−1.00; −0.21]). In snails from the open habitat however, there was no such difference between morphs *Δ*_[5bands–0bands]_ = 0.17 SDs [−0.19; 0.59]). This was due to unbanded snails being less mobile when coming from the open habitat than when coming from the shaded population *Δ*_[shaded–open]_ = 0.71 SDs [0.31; 1.08]), while there was no clear between-population difference for five-banded snails (*Δ*_[shaded–open]_ = −0.08 SDs [−0.47; 0.32])(**Table 1, Fig. 4).** In any case, there was no detectable link between movement and shell size **(Table 1**).

**Figure 4.**
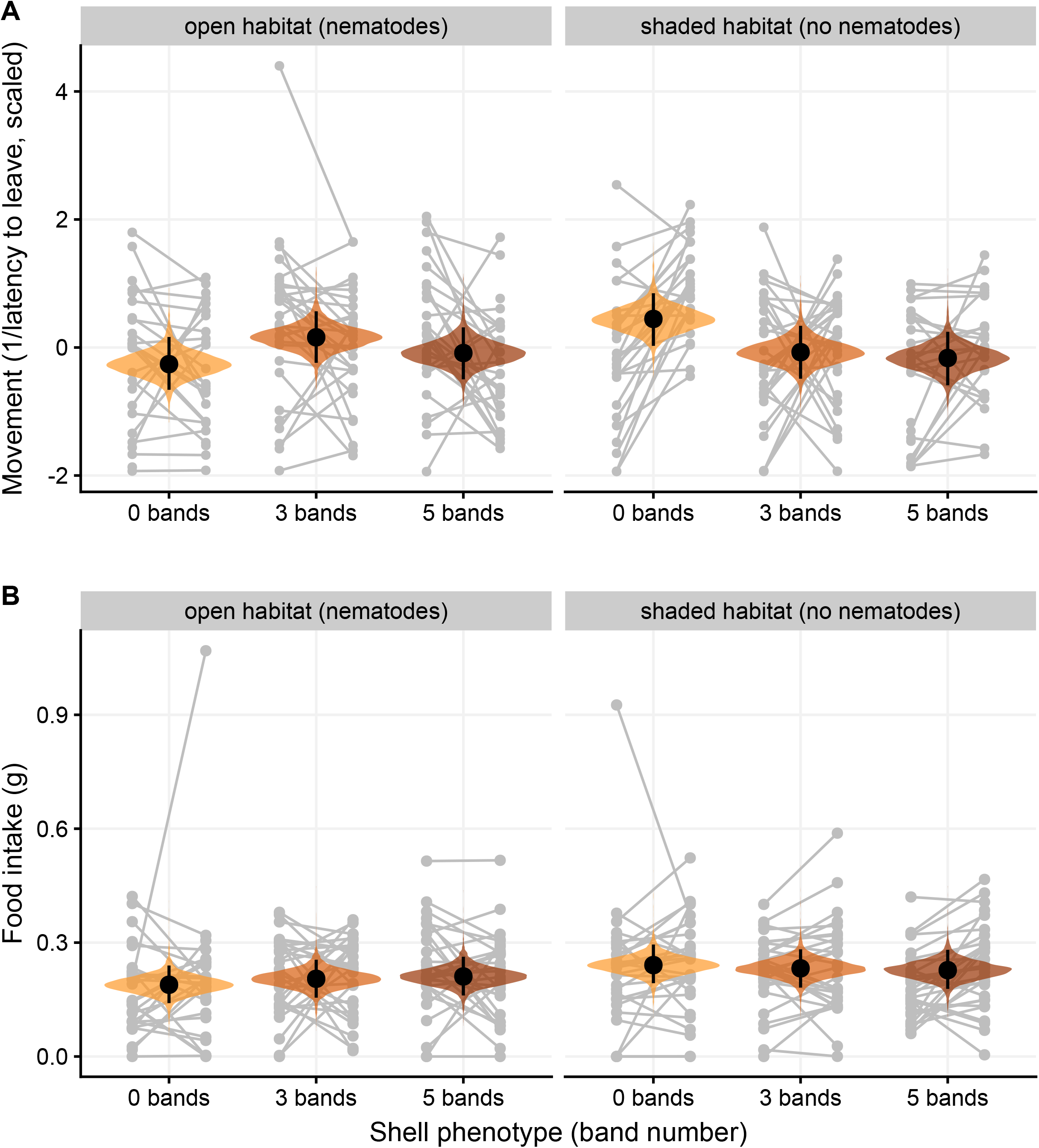
Behaviours as a function of morph and population of origin. (A) Movement metric (1/latency to move beyond 10 cm, centered and scaled to unit 1SD); (B) Food intake. Grey dots are observed values, displayed along the posterior distributions of the average abundances based on the model described in **Table 1** (black dots and segments are the means and 95% CI of the posteriors). Observed values from the same individual are connected by a line. Posterior predictions are based on fixed effects only, and the effects of shell size are averaged out. See **Supporting Information S3** for pairwise comparisons.

Larger snails tended to eat more (**Table 1**). Source population had no effect on food intake if snails were three- (*Δ*_[shaded–open]_ = 0.03 g [−0.02; 0.08], **Fig. 4**) or five-banded (*Δ*_[shaded–open]_ = 0.02 g [−0.03; 0.07], Fig. 4), but similarly to movement, unbanded snails coming from the shaded population ate more on average than their counterparts from the open habitat (*Δ*_[shaded–open]_ = 0.05 g [0.00; 0.10], Fig. 4).

Looking at the individual-level correlation matrix between traits (i.e. once fixed effects of shell morph, size and population of origin are accounted for), food intake and nematode abundance are negatively correlated, while all other correlations (including between nematode and trematode abundance) are not markedly different from zero **(Table 2).** The food-nematode correlation persists if we refit the nematode model to include data from the shaded population **(−0.58 [−0.87; −0.29]).**

**Table 2.** Individual-level correlations between parasite abundances and behavioural traits. Correlations involving nematode abundance are only based on individuals from the open population (see Methods); results are not qualitatively different if the model is refitted to include all observations (see **Data availability).** Correlations that are different from 0 (based on 95% intervals) are in bold.

|  | Nematode abundance | Trematode abundance | Movement |
| --- | --- | --- | --- |
| **Trematode abundance** | 0.01 [-0.35; 0.37] | -- | -- |
| **Movement** | -0.24 [-0.73; 0.25] | -0.17 [-0.68; 0.28] | -- |
| **Food intake** | **-0.60 [-0.87; -0.30]** | -0.12 [-0.46; 0.19] | 0.24 [-0.23; 0.71] |

## Discussion

To understand how individual variation in behaviour and infection interact, we studied movement, food intake and natural parasite infection across two populations and three morphs of the land snail *Cepaea nemoralis. We* found evidence of morph-dependent responses to parasites, as well as links between parasite infection and behaviour. We additionally present evidence of land snails encapsulating parasitic arthropods (mites) in their shell.

Snails of the darker morphs were smaller in the shaded population (Fig. 1). While the two habitats differ along various abiotic dimensions (calcium availability in substrate, microclimate…, see **Methods),** their documented effect on shell size are highly species- and study-dependent, and there is no consistent evidence for or against their role in driving the differences we observe (Goodfriend, 1986). Differences in parasite infection between the two habitats might also influence body size, for instance if high parasite risk drives resource allocation away from growth and towards immune defence (Rantala & Roff, 2005). In any case, our models conditioned on size, meaning that any effect of morph/population we discuss below is “size being equal”. In addition, we did not find any effect of size itself on parasites or movement **(Table 1**), meaning that the effects of morph/population we found are likely not mediated by size differences.

One snail was found with live *Riccardoella* mites, hemolymph feeders with negative effects on mollusc life-history and behavior (e.g. Schüpbach & Baur, 2008a). They are transmitted horizontally during mating, but may find new hosts by leaving their host and following snail mucus trails (Schüpbach & Baur, 2008b). However, their limited prevalence in our sample prevented us from investigating their effects. Intriguingly, we found encapsulated mites in three other snails **(Fig. 2).** Many land snails can defend themselves against nematode infection by trapping them in their shell (Rae, 2017), including *Cepaea nemoralis* (this study, Williams & Rae, 2016; Rae, 2017). Based on extensive surveys, Rae (2017) wrote that encapsulation in the shell is a specific response against nematode infection; our observations might contradict this affirmation. *Riccardoella* infection is highly variable across populations or species, with reported prevalences of 0% or near 0% being frequently observed (this study; Russo & Madec, 2013; Baur & Baur, 2005; Segade *et al*., 2013), which may explain why encapsulated mites were not previously found. We need further studies to determine with certainty whether shell encapsulation is a more general response to metazoan parasites or if it is a nematode-specific response, as posited by Rae (2017), but occasionally and anecdotally catches other foreign bodies.

A majority of snails in the open population were hosting nematodes, most of which were likely Rhabditidae based on size and organ location (Morand *et al*., 2004; Wilson, 2012). Interactions between Rhabditidae and terrestrial molluscs are complex, including phoresy/endozoochory, facultative parasitism, necromeny (feeding and reproducing on the host’s body after its death), and obligate larval parasites that actively infect snails through e.g. the lung cavity (Morand *et al*., 2004; Pieterse *et al*., 2017; Sudhaus, 2018). Delimiting these interaction types is complicated because even when they have no effects on adults, these nematodes can have dramatically negative effects on juveniles (Grannell, Cutler & Rae, 2021). By contrast to the open population, we found no live nematodes in any snail from the shaded population **(Fig. 3;** note that our methods were not designed to detect nematodes embedded in e.g. snail foot tissue). Thanks to nematodes trapped in shells **(Fig. 3),** which integrate infections over time, we know parasitic nematodes are not truly absent from the shaded population; they were simply not infecting snails at the time of sampling. The “shaded” woodlot habitat is sheltered from the sun, more humid and likely colder than the open one. Infections are often dependent on abiotic parameters, including temperature and humidity, leading to seasonality (Altizer *et al*., 2006). Differences in nematode infection between the habitats may then be explained, at least partly, by differences in microclimate. First, the different microclimates experienced by the two populations may favour different nematode species within the soil community, differing in their tendency to parasitism. Second, even if nematode communities are similar, conditions favourable to nematode activity, development or infection may be reached only later in the year in the shaded habitat. For instance, *Deroceras reticulatum* slugs were more negatively impacted by *Phasmarhabditis hermaphrodita* infection when conditions were artificially warmed during winter (El-Danasoury & Iglesias-Piñeiro, 2017), indicating that warmer conditions may favour nematode activity. In any case, these results mean that any between-population differences in snail behaviour in the present study should be interpreted not only in relation with microhabitat and microclimate differences, but also in terms of average infection levels.

In the open population where active nematodes were actually found, morph differences in infection seem to validate our initial hypothesis of increased immune defence in darker snails **(Fig. 3).** By contrast, no significant morph difference in immune defence was found in the related *Cepaea hortensis* (Scheil *et al*., 2013, 2014), but their endpoints were different (phenoloxidase activity and mortality). In addition, our behavioural results (discussed below) are also consistent with the hypothesis of morph-dependent immunity. These results are interesting in the light of recent evidence that melanins may not be the pigments forming the dark bands in *C. nemoralis* (Affenzeller *et al*., 2020). Indeed, our initial hypothesis when designing this experiment (in 2019) was based on the oft-seen association between immunity and melanin (San-Jose & Roulin, 2018), including in snails (Coaglio *et al*., 2018), although other pigments may also have links to immunity (Tan *et al*., 2020). While it is still likely that melanin-associated pathways play a role in immune defence in *Cepaea* (Scheil *et al*., 2014), it is probably not as simple as “banded snails constitutively produce more melanin, and thus are better protected”. More studies are needed to understand the mechanisms linking shell morph to nematode resistance.

Snails from all morphs and both populations were infected by *Brachylaima* trematodes **(Fig. 3).** We previously documented *Brachylaima* infecting *Cornu aspersum* in the same area (Gérard *et al*., 2020); finding these parasites a year later and in *C. nemoralis* may indicate that they are well-implanted in local communities. Contrary to nematodes, we found no evidence that morph or population influenced trematode abundance **(Fig. 3).** This is consistent with another study analyzing both *C. nemoralis* and *C. hortensis* (Żbikowska *et al*., 2020). We must acknowledge however that snail infections by *Brachylaima* vary seasonally, and that spring may not be the infection peak (Gracenea & Gállego, 2017). Longitudinal studies, possibly combined with experimental infections or cures, may help confirm whether morphs respond differently to trematode risk.

Nematode and trematode numbers were uncorrelated **(Table 2).** Terrestrial gastropods are commonly infected by both nematodes and trematodes at the same time (e.g. Segade *et al*., 2013; Antzée-Hyllseth *et al*., 2020). Our results extend these previous studies, and suggest that despite sharing a host, nematode and trematode niches are sufficiently distinct within snails to limit spatial and trophic competition. We reiterate however that studies more expansive in space and time, including seasons and/or populations with higher prevalence and abundance of trematodes and nematodes, are needed to confirm this. Combined with finer-grained studies (e.g. at the parasite species level; Morand *et al*., 2004), this would allow us to understand better how parasites with similar or different niches within snails interact.

We found that movement behaviour in *C. nemoralis* was morph-dependent, but that this dependency varied between populations: five-banded morphs were less mobile, but contrary to Dahirel et al. (2021), this was only the case in the shaded population **(Fig. 4).** In the open population, snail morphs are mobile at similarly low levels: five-banded snails do not change behaviour between the two populations, while unbanded snails go down to five-banded snail levels of mobility **(Fig. 4).** One key difference between the present study and Dahirel et al. (2021) is the collection date. While here snails were collected several weeks after the onset of spring activity and kept active throughout, in Dahirel et al. (2021) they were collected in late fall and stored at cold temperatures inducing dormancy for months before being tested. This difference may be especially relevant: winter or summer dormancy can lead to sharp decreases in parasite infection in several parasite-snail systems (Solomon, Paperna & Matkovics, 1996; Haeussler *et al*., 2012; Segade *et al*., 2013). We can thus reconcile the behavioral results from Dahirel et al. (2021) from those of the present study if we assume that in the former experiment, hibernation reduced parasite load across all morphs and populations, and that in the present study, lower temperatures in the shaded habitat led to longer hibernation. We can then interpret population differences in the present study in the light of their different infection patterns. Five-banded snails’ behaviour is almost unchanged between the two populations, while unbanded snails from the “infected” population are slower than in the nematode-free population **(Fig. 4).** This is again consistent with the idea that unbanded snails are more vulnerable to parasites than banded ones. The question then becomes: is this reduction in activity in infected or at-risk unbanded snails simply the direct consequence of the costs of infection, or does it reflect something more? For instance, infective nematodes may be able to follow mucus trails from active snails (Andrus *et al*., 2020). Reducing activity and the associated mucus production may then be protective for snails, by limiting the risk of further infections. Experimental manipulations of both snail infection and parasite risk in the environment may help disentangle these two responses. Future studies including species-level identification of nematodes will also be needed, to determine whether nematodes with more negative impacts also trigger stronger behavioural responses, or if snails respond indiscriminately to nematode abundance.

Food intake was influenced by nematode infection: highly parasitized snails ate less **(Table 2).** Feeding reduction in nematode-infected individuals has been documented in other terrestrial molluscs (Antzée-Hyllseth *et al*., 2020; Saeedizadeh & Niasti, 2020) and is an indication that parasite infection is indeed costly for snails even if they are non-lethal, at least in adults (Wilson *et al*., 2000; Williams & Rae, 2016; Grannell *et al*., 2021). By contrast, trematode infection had no effect on behaviour, including feeding behaviour **(Table 2).** We however only found snails infected by *Brachylaima* metacercariae. At the sporocyst stage, *Brachylaima* parasitizes the digestive gland (e.g. Gérard *et al*., 2020), which would likely have a stronger effect on snail behaviour. Indeed, sporocysts can cause high damage, direct and indirect, to that key organ for digestion and for energy storage (Segade *et al*., 2011; Żbikowska *et al*., 2020).

The fact that snail response to parasites seems morph-dependent for nematodes, but not trematodes, has interesting implications for host-parasite and snail-predator interactions in a changing world. Trematodes infecting land snails are trophically transmitted to vertebrate definitive hosts including rodents and birds (Gérard *et al*., 2020; Sitko & Heneberg, 2020). Visual selection by thrushes (*Turdus philomelos*) is held as one of the key explanations for the maintenance of *Cepaea* shell polymorphism (Jones *et al*., 1977), and predation by rodents may also be morph-dependent (Rosin *et al*., 2011; Rosin *et al*., 2013). If confirmed, our results suggest that morph-dependent predation confers no additional trematode-avoiding advantage.

In addition, snail morphs differ in their thermal responses: lighter-coloured morphs are better adapted to heat, becoming more frequent with global warming or urbanization (Jones *et al*., 1977; Silvertown *et al*., 2011; Ożgo *et al*., 2017; Kerstes *et al*., 2019). If lighter morphs are more vulnerable to infection by nematodes, then we need to understand how snail immunity will interact with thermal tolerance to predict how snail and nematode dynamics will be altered in response to climate change (El-Danasoury & Iglesias-Piñeiro, 2017; Aleuy & Kutz, 2020).

## Acknowledgements

We thank participants to the 4th French Conference on Animal Ecophysiology (CEPA4, Rennes, 2019) for interesting discussions. Funding for MD’s position was obtained by Elodie Vercken (see Funding).

## Data availability

Data and R scripts to reproduce all analyses presented here are on Github (https://github.com/mdahirel/cepaea-parasite-behaviour-2019) and archived in Zenodo (DOI: https://doi.org/10.5281/zenodo.6327265). Representative images of parasites trapped in shells are also archived in Zenodo (DOI: https://doi.org/10.5281/zenodo.6326305).

## References

Affenzeller, S., Wolkenstein, K., Frauendorf, H. & Jackson, D.J. (2020). Challenging the concept that eumelanin is the polymorphic brown banded pigment in *Cepaea nemoralis*. Sci. Rep. 10, 2442.

Aleuy, O.A. & Kutz, S. (2020). Adaptations, life-history traits and ecological mechanisms of parasites to survive extremes and environmental unpredictability in the face of climate change. Int. J. Parasitol. Parasites Wildl. 12, 308–317.

Altizer, S., Dobson, A., Hosseini, P., Hudson, P., Pascual, M. & Rohani, P. (2006). Seasonality and the dynamics of infectious diseases. Ecol. Lett. 9, 467–484.

Andrus, P., Ingle, O., Coleman, T. & Rae, R. (2020). Gastropod parasitic nematodes *(Phasmarhabditis* sp.) are attracted to hyaluronic acid in snail mucus by cGMP signalling. J. Helminthol. 94, e9.

Antzée-Hyllseth, H., Trandem, N., Torp, T. & Haukeland, S. (2020). Prevalence and parasite load of nematodes and trematodes in an invasive slug and its susceptibility to a slug parasitic nematode compared to native gastropods. J. Invertebr. Pathol. 173, 107372.

Barker, G.M. (Ed.). (2004). Natural enemies of terrestrial molluscs. Wallingford, UK: CABI.

Baur, A. & Baur, B. (2005). Interpopulation variation in the prevalence and intensity of parasitic mite infection in the land snail *Arianta arbustorum*. Invertebr. Biol. 124, 194–201.

Binning, S.A., Shaw, A.K. & Roche, D.G. (2017). Parasites and host performance: incorporating infection into our understanding of animal movement. Integr. Comp. Biol. 57, 267–280.

BRGM. (2005). Geological map at 1:50 000 (BD-CHARM-50), department: Deux-Sèvres (79, France).

Bürkner, P.-C. (2017). brms: an R package for Bayesian multilevel models using Stan. J. Stat. Softw. 80, 1–28.

Butcher, A.R. (2016). Children, snails and worms: the *Brachylaima cribbi* story. Microbiol. Aust. 37, 30–33.

Cain, A.J. & Sheppard, P.M. (1954). Natural selection in *Cepaea*. Genetics 39, 89–116.

Carpenter, B., Gelman, A., Hoffman, M.D., Lee, D., Goodrich, B., Betancourt, M., Brubaker, M., Guo, J., Li, P. & Riddell, A. (2017). Stan: a probabilistic programming language. J. Stat. Softw. 76, 1–32.

Coaglio, A.L., Ferreira, M.A.N.D., dos Santos Lima, W. & de Jesus Pereira, C.A. (2018). Identification of a phenoloxidase-and melanin-dependent defence mechanism in *Achatina fulica* infected with *Angiostrongylus vasorum*. Parasites Vectors 11, 113.

Dahirel, M., Gaudu, V. & Ansart, A. (2021). Boldness and exploration vary between shell morphs but not environmental contexts in the snail *Cepaea nemoralis*. Ethology 13.

Davies, M.S. & Knowles, A.J. (2001). Effects of trematode parasitism on the behaviour and ecology of a common marine snail *(Littorina littorea* (L.)). J. Exp. Mar. Biol. Ecol. 260, 155–167.

Debeffe, L., Morellet, N., Verheyden-Tixier, H., Hoste, H., Gaillard, J.-M., Cargnelutti, B., Picot, D., Sevila, J. & Hewison, A.J.M. (2014). Parasite abundance contributes to condition-dependent dispersal in a wild population of large herbivore. Oikos 123, 1121–1125.

Douma, J.C. & Weedon, J.T. (2019). Analysing continuous proportions in ecology and evolution: a practical introduction to beta and Dirichlet regression. Methods Ecol. Evol. 10, 1412–1430.

Dupuis, J., Cariou, E., Coirier, B. & Ducloux, J. (1983). Notice explicative, Carte géologigue de la France à 1/50 000,feuille Niort (610). Orléans, France: BRGM.

El-Danasoury, H. & Iglesias-Piñeiro, J. (2017). Performance of the slug parasitic nematode *Phasmarhabditis hermaphrodita* under predicted conditions of winter warming. J. Pestic. Science 42, 62–66.

Ezenwa, V.O., Archie, E.A., Craft, M.E., Hawley, D.M., Martin, L.B., Moore, J. & White, L. (2016). Host behaviour–parasite feedback: an essential link between animal behaviour and disease ecology. Proc. Royal Soc. B 283, 20153078.

Fain, A. (2004). Mites (Acari) parasitic and predaceous on terrestrial gastropods. In Natural enemies of terrestrial molluscs: 505–524. Barker, G.M. (Ed.). Wallingford, UK: CABI.

Furuta, E. & Yamaguchi, K. (2001). Haemolymph: blood cell morphology and function. In The biology of terrestrial molluscs: 289–306. Barker, G.M. (Ed.). Wallingford, UK: CABI.

Gabry, J., Simpson, D., Vehtari, A., Betancourt, M. & Gelman, A. (2019). Visualization in Bayesian workflow. J. Royal Stat. Soc. A 182, 389–402.

Gérard, C., Ansart, A., Decanter, N., Martin, M.-C. & Dahirel, M. (2020). *Brachylaima* spp. (Trematoda) parasitizing *Cornu aspersum* (Gastropoda) in France with potential risk of human consumption. Parasite 27, 15.

Giannelli, A., Cantacessi, C., Colella, V., Dantas-Torres, F. & Otranto, D. (2016). Gastropod-borne helminths: a look at the snail–parasite interplay. Trends Parasitol. 32, 255–264.

González-Santoyo, I. & Córdoba-Aguilar, A. (2012). Phenoloxidase: a key component of the insect immune system. Entomol. Exp. Appl. 142, 1–16.

Goodfriend, G.A. (1986). Variation in land-snail shell form and size and its causes: a review. Syst. Zool. 35, 204–223.

Gracenea, M. & Gállego, L. (2017). Brachylaimiasis: *Brachylaima* spp. (Digenea: Brachylaimidae) metacercariae parasitizing the edible snail *Cornu aspersum* (Helicidae) in Spanish public marketplaces and health-associated risk factors. J. Parasitol. 103, 440–450.

Grannell, A., Cutler, J. & Rae, R. (2021). Size-susceptibility of *Cornu aspersum* exposed to the malacopathogenic nematodes *Phasmarhabditis hermaphrodita* and *P. californica*. Biocontrol Sci. Technol. 31, 1149–1160.

Haeussler, E.M., Pizá, J., Schmera, D. & Baur, B. (2012). Intensity of parasitic mite infection decreases with hibernation duration of the host snail. Parasitology 139, 1038–1044.

Harrison, X.A. (2014). Using observation-level random effects to model overdispersion in count data in ecology and evolution. PeerJ 2, e616.

Jones, J.S., Leith, B.H. & Rawlings, P. (1977). Polymorphism in *Cepaea*: a problem with too many solutions? Annu. Rev. Ecol. Evol. Syst. 8, 109–143.

Kavaliers, M. (1992). Opioid systems, behavioral thermoregulation and shell polymorphism in the land snail, *Cepaea nemoralis*. J. Comp. Physiol. B 162, 172–178.

Kay, M. (2019). tidybayes: tidy data and geoms for Bayesian models. https://doi.org/10.5281/zenodo.1308151

Kerstes, N.A.G., Breeschoten, T., Kalkman, V.J. & Schilthuizen, M. (2019). Snail shell colour evolution in urban heat islands detected via citizen science. Commun. Biol. 2, 264.

Köhler, H.-R., Capowiez, Y., Mazzia, C., Eckstein, H., Kaczmarek, N., Bilton, M.C., Burmester, J.K.Y., Capowiez, L., Chueca, L.J., Favilli, L., Gomila, J.F., Manganelli, G., Mazzuca, S., Moreno-Rueda, G., Peschke, K., Piro, A., Cardona, J.Q., Sawallich, L., Staikou, A.E., Thomassen, H.A. & Triebskorn, R. (2021). Experimental simulation of environmental warming selects against pigmented morphs of land snails. Ecol. Evol. 11, 1111–1130.

Lamotte, M. (1959). Polymorphism of natural populations of *Cepaea nemoralis*. Cold Spring Harb. Symp. Quant. Biol. 24, 65–86.

McElreath, R. (2020). Statistical rethinking: a Bayesian course with examples in R and Stan. 2nd edition. Boca Raton, USA: Chapman and Hall/CRC.

McElroy, E.J. & de Buron, I. (2014). Host performance as a target of manipulation by parasites: a meta-analysis. J. Parasitol. 100, 399–410.

Meerburg, B.G., Singleton, G.R. & Kijlstra, A. (2009). Rodent-borne diseases and their risks for public health. Crit. Rev. Microbiol. 35, 221–270.

Morand, S., Wilson, M.J. & Glen, D.M. (2004). Nematodes (Nematoda) parasitic in terrestrial molluscs. In Natural enemies of terrestrial molluscs: 525–557. Barker, G.M. (Ed.). Wallingford, UK: CABI.

Morley, N.J. & Lewis, J.W. (2008). The influence of climatic conditions on long-term changes in the helminth fauna of terrestrial molluscs and the implications for parasite transmission in southern England. J. Helminthol. 82, 325–335.

Nakagawa, S. & Schielzeth, H. (2010). Repeatability for Gaussian and non-Gaussian data: a practical guide for biologists. Biol. Rev. 85, 935–956.

O’Dwyer, K., Kamiya, T. & Poulin, R. (2014). Altered microhabitat use and movement of littorinid gastropods: the effects of parasites. Mar. Biol. 161, 437–445.

Ożgo, M. (2009). Current problems in the research of *Cepaea* polymorphism. Folia Malacol. 16, 55–60.

Ożgo, M., Liew, T.-S., Webster, N.B. & Schilthuizen, M. (2017). Inferring microevolution from museum collections and resampling: lessons learned from *Cepaea*. PeerJ 5, e3938.

Pedersen, T.L. (2020). patchwork: the composer of plots. https://CRAN.R-project.org/package=patchwork

Pieterse, A., Malan, A.P. & Ross, J.L. (2017). Nematodes that associate with terrestrial molluscs as definitive hosts, including *Phasmarhabditis hermaphrodita* (Rhabditida: Rhabditidae) and its development as a biological molluscicide. J. Helminthol. 91, 517–527.

R Core Team. (2021). R: a language and environment for statistical computing. Vienna, Austria: R Foundation for Statistical Computing.

Rae, R. (2017). The gastropod shell has been co-opted to kill parasitic nematodes. Sci. Rep. 7, 4745.

Rantala, M.J. & Roff, D.A. (2005). An analysis of trade-offs in immune function, body size and development time in the Mediterranean Field Cricket, *Gryllus bimaculatus*. Funct. Ecol. 19, 323–330.

Réale, D., Garant, D., Humphries, M.M., Bergeron, P., Careau, V. & Montiglio, P.-O. (2010). Personality and the emergence of the pace-of-life syndrome concept at the population level. Phil. Trans. R. Soc. B 365, 4051–4063.

Rosin, Z.M., Kobak, J., Lesicki, A. & Tryjanowski, P. (2013). Differential shell strength of *Cepaea nemoralis* colour morphs—implications for their anti-predator defence. Naturwissenschaften 100, 843–851.

Rosin, Z.M., Olborska, P., Surmacki, A. & Tryjanowski, P. (2011). Differences in predatory pressure on terrestrial snails by birds and mammals. J. Biosci. 36, 691–699.

Royauté, R., Berdal, M.A., Garrison, C.R. & Dochtermann, N.A. (2018). Paceless life? A meta-analysis of the pace-of-life syndrome hypothesis. Behav. Ecol. Sociobiol. 72, 64.

Russo, J. & Madec, L. (2013). Linking immune patterns and life history shows two distinct defense strategies in land snails (Gastropoda, Pulmonata). Physiol. Biochem. Zool. 86, 193–204.

Saeedizadeh, A. & Niasti, F. (2020). Response of grey slug to entomopathogenic nematodes. Bragantia 79, 447–456.

San-Jose, L.M. & Roulin, A. (2018). Toward understanding the repeated occurrence of associations between melanin-based coloration and multiple phenotypes. Am. Nat. 192, 111–130.

Scheil, A.E., Hilsmann, S., Triebskorn, R. & Köhler, H.-R. (2013). Shell colour polymorphism, injuries and immune defense in three helicid snail species, *Cepaea hortensis, Theba pisana* and *Cornu aspersum maximum*. Results Immunol. 3, 73–78.

Scheil, A.E., Hilsmann, S., Triebskorn, R. & Köhler, H.-R. (2014). Shell colouration and parasite tolerance in two helicoid snail species. J. Invertebr. Pathol. 117, 1–8.

Schilthuizen, M. (2013). Rapid, habitat-related evolution of land snail colour morphs on reclaimed land. Heredity 110, 247–252.

Schüpbach, H.U. & Baur, B. (2008a). Parasitic mites influence fitness components of their host, the land snail *Arianta arbustorum*. Invertebr. Biol. 127, 350–356.

Schüpbach, H.U. & Baur, B. (2008b). Experimental evidence for a new transmission route in a parasitic mite and its mucus-dependent orientation towards the host snail. Parasitology 135, 1679–1684.

Segade, P., Crespo, C., García, N., García-Estévez, J.M., Arias, C. & Iglesias, R. (2011). *Brachylaima aspersae* n. sp. (Digenea: Brachylaimidae) infecting farmed snails in NW Spain: Morphology, life cycle, pathology, and implications for heliciculture. Vet. Parasitol. 175, 273–286.

Segade, P., García-Estévez, J., Arias, C. & Iglesias, R. (2013). Parasitic infections in mixed system-based heliciculture farms: dynamics and key epidemiological factors. Parasitology 140, 482–497.

Silvertown, J., Cook, L., Cameron, R., Dodd, M., McConway, K., Worthington, J., Skelton, P., Anton, C., Bossdorf, O., Baur, B., Schilthuizen, M., Fontaine, B., Sattmann, H., Bertorelle, G., Correia, M., Oliveira, C., Pokryszko, B., Ożgo, M., Stalažs, A., Gill, E., Rammul, Ü., Sólymos, P., Féher, Z. & Juan, X. (2011). Citizen science reveals unexpected continental-scale evolutionary change in a model organism. PLOS ONE 6, e18927.

Sitko, J. & Heneberg, P. (2020). Emerging helminthiases of song thrush *(Turdus philomelos)* in Central Europe. Parasitol Res. 119, 4123–4134.

Solomon, A., Paperna, I. & Matkovics, A. (1996). The influence of aestivation in land snails on the larval development of *Muellerius* cf. *capillaris* (Metastrongyloidea: Protostrongylidae). Int. J. Parasitol. 26, 363–367.

Sudhaus, W. (2018). Dispersion of nematodes (Rhabditida) in the guts of slugs and snails. Soil Org. 90, 101–114.

Surmacki, A., Ożarowska-Nowicka, A. & Rosin, Z.M. (2013). Color polymorphism in a land snail *Cepaea nemoralis* (Pulmonata: Helicidae) as viewed by potential avian predators. Naturwissenschaften 100, 533–540.

Tan, K., Zhang, H., Lim, L.-S., Ma, H., Li, S. & Zheng, H. (2020). Roles of carotenoids in invertebrate immunology. Front. Immunol. 10.

Vehtari, A., Gelman, A., Simpson, D., Carpenter, B. & Bürkner, P.-C. (2021). Rank-normalization, folding, and localization: an improved 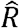 for assessing convergence of MCMC (with discussion). Bayesian Anal. 16, 667–718.

Wickham, H., Averick, M., Bryan, J., Chang, W., McGowan, L., François, R., Grolemund, G., Hayes, A., Henry, L., Hester, J., Kuhn, M., Pedersen, T., Miller, E., Bache, S., Müller, K., Ooms, J., Robinson, D., Seidel, D., Spinu, V., Takahashi, K., Vaughan, D., Wilke, C., Woo, K. & Yutani, H. (2019). Welcome to the Tidyverse. J. Open Source Softw. 4, 1686.

Williams, A. & Rae, R. (2016). *Cepaea nemoralis* (Linnaeus, 1758) uses its shell as a defence mechanism to trap and kill parasitic nematodes. J. Molluscan Stud. 82, 349–350.

Williams, S.T. (2017). Molluscan shell colour. Biol. Rev. 92, 1039–1058.

Wilson, M.J. (2012). Pathogens and parasites of terrestrial molluscs. In Manual of techniques in invertebrate pathology: 427–439. Lacey, L.A. (Ed.). Oxford⍰ New York: Academic Press.

Wilson, M.J., Hughes, L.A., Hamacher, G.M. & Glen, D.M. (2000). Effects of *Phasmarhabditis hermaphrodita* on non-target molluscs. Pest. Manag. Sci. 56, 711–716.

Wynne, R., Morris, A. & Rae, R. (2016). Behavioural avoidance by slugs and snails of the parasitic nematode *Phasmarhabditis hermaphrodita*. Biocontrol Sci. Technol. 26, 1129–1138.

Żbikowska, E., Marszewska, A., Cichy, A., Templin, J., Smorąg, A. & Strzała, T. (2020). *Cepaea* spp. as a source of *Brachylaima mesostoma* (Digenea: Brachylaimidae) and *Brachylecithum* sp. (Digenea: Dicrocoeliidae) larvae in Poland. Parasitol. Res. 119, 145–152.

