## Supplementary Material for "Morph-dependent nematode infection and its association with host movement in the land snail *Cepaea nemoralis* (Mollusca, Gastropoda)"

### Supplementary Material for "Morph-dependent effect of nematode infection on host movement in the land snail *Cepaea nemoralis* (Mollusca, Gastropoda)"

Maxime Dahirel, Marine Proux, Claudia Gérard, Armelle Ansart

#### S1 - Detailed model descriptions

In both models described below, we used weakly informative priors inspired by McElreath (2020). We used $\text{Normal}(0, 1)$ priors for the fixed effects $\beta$, and $\text{Half} - \text{Normal}(0, 1)$ priors for all standard deviations (random effect standard deviations $\sigma_\alpha$ and $\sigma_\gamma$ as well as residual standard deviations $\sigma_r$). We used a $\text{LKJ}(2)$ prior for the individual-level correlation matrix in the multivariate model.

##### Body size (shell diameter) model

After centering and standardizing to unit 1SD, shell diameters $D_{o,m,i}$ with $o$ the population of origin (coded as a binary variable with 0 for the sun-exposed population and 1 for the shaded population), $m$ the shell morph and $i$ the individual, can be described by the following linear model:

$$D_{o,m,i} \sim \text{Normal}(\mu_{o,m}, \sigma_{r[D]}),$$
$$\mu_{o,m} = \beta_{0[D,m]} + \beta_{1[D,m]} \times o.$$

In this model, the intercepts $\beta_0$ are the morph-specific intercepts/mean sizes in the sun-exposed habitat, $\beta_1$ the morph-specific effect of the shaded habitat, and $\sigma_{r[D]}$ the residual variation.

##### Multivariate model of parasite abundance and snail behaviours

The multivariate model used to analyse parasite abundances, movement behaviour, food intake and their individual-level correlations can be written as follows.

Let $N_{o,m,s,i}$ and $T_{o,m,s,i}$ be the numbers of live nematodes and *Brachylaima* trematodes (metacercariae) found in snail $i$ which was tested during session $s$, and $F_{o,m,s,i,t}$ and $M_{o,m,s,i,t}$ be the quantity of food ingested by snail $i$ (centered and scaled to 1SD), and its movement activity during trial $t$ (also centered and scaled to 1SD), respectively. We then have:

$$\begin{cases} N_{o,m,s,i} \sim \text{Poisson}(\lambda_{m,s,i[N]}), & \text{if } o = 0 \text{ (sun-exposed population)} \\ N_{o,m,s,i} = 0, & \text{if } o = 1 \text{ (shaded population)}, \end{cases}$$

$$T_{o,m,s,i} \sim \text{Poisson}(\lambda_{o,m,s,i[T]}),$$
$$F_{o,m,s,i,t} \sim \text{Normal}(\mu_{o,m,s,i[F]}, \sigma_{r[F]}),$$
$$M_{o,m,s,i,t} \sim \text{Normal}(\mu_{o,m,s,i[M]}, \sigma_{r[M]}),$$
$$\lambda_{m,s,i[N]} = \beta_{0[N,m]} + \beta_{2[N]} \times D_i + \alpha_{s[N]} + \gamma_{i[N]},$$

$$\lambda_{o,m,s,i[T]} = \beta_{0[T,m]} + \beta_{1[T,m]} \times o + \beta_{2[T]} \times D_i + \alpha_{s[T]} + \gamma_{i[T]},$$

$$\mu_{o,m,s,i[F]} = \beta_{0[F,m]} + \beta_{1[F,m]} \times o + \beta_{2[F]} \times D_i + \alpha_{s[F]} + \gamma_{i[F]},$$

$$\mu_{o,m,s,i[M]} = \beta_{0[M,m]} + \beta_{1[M,m]} \times o + \beta_{2[M]} \times D_i + \alpha_{s[M]} + \gamma_{i[M]},$$

where the $\alpha$ refer to the experimental session random effects and $\gamma$ the individual random effects. These random effects are distributed as follows:

$$\alpha_{s[N]} \sim \text{Normal}(0, \sigma_{\alpha[N]}),$$

$$\alpha_{s[T]} \sim \text{Normal}(0, \sigma_{\alpha[T]}),$$

$$\alpha_{s[F]} \sim \text{Normal}(0, \sigma_{\alpha[F]}),$$

$$\alpha_{s[M]} \sim \text{Normal}(0, \sigma_{\alpha[M]}),$$

$$\begin{bmatrix} \gamma_{i[N]} \\ \gamma_{i[T]} \\ \gamma_{i[F]} \\ \gamma_{i[M]} \end{bmatrix} \sim \text{MVNormal}\left( \begin{bmatrix} 0 \\ 0 \\ 0 \\ 0 \end{bmatrix}, \boldsymbol{\Omega} \right),$$

, where $\boldsymbol{\Omega}$ is the individual-level covariance matrix, which can be decomposed into its constituent standard deviations and correlation matrix $\boldsymbol{R}$ in this way:

$$\boldsymbol{\Omega} = \begin{bmatrix} \sigma_{\gamma[N]} & 0 & 0 & 0 \\ 0 & \sigma_{\gamma[T]} & 0 & 0 \\ 0 & 0 & \sigma_{\gamma[F]} & 0 \\ 0 & 0 & 0 & \sigma_{\gamma[M]} \end{bmatrix} \boldsymbol{R} \begin{bmatrix} \sigma_{\gamma[N]} & 0 & 0 & 0 \\ 0 & \sigma_{\gamma[T]} & 0 & 0 \\ 0 & 0 & \sigma_{\gamma[F]} & 0 \\ 0 & 0 & 0 & \sigma_{\gamma[M]} \end{bmatrix}.$$

### S2 - Summary statistics on parasite infections and encapsulations

In the following tables, we present detailed descriptive information about the prevalence of each parasite (proportion of snails infected) and the infection intensity (number of parasites per *actually infected* snail). Parasite abundance, which we used as a synthetic measure of infection in the main text, is the mean number of parasites per snail *all snails included*; mean parasite abundance is equal to prevalence $\times$ mean intensity. See Bush *et al.* (1997) for details on definitions.

For trematode and nematode data, the provided 95% intervals are Highest Density intervals based on Bernoulli (prevalence) and negative binomial (intensity) GLMs including shell morph, population and their interaction as fixed effects. The intensity models are fitted on (number of parasites - 1), to account for the zero truncation inherent to intensity. Priors are as in **S1**, with the addition of a $\text{Half} - \text{Normal}(0, 1)$ prior on the inverse of the negative binomial shape parameter, inspired by the authors of the Stan wiki: https://github.com/stan-dev/stan/wiki/Prior-Choice-Recommendations.

Note that we do not provide 95% intervals for mites data, given how rare infections are in our dataset. We also do not provide either intensity information or 95% intervals for encapsulated parasites, for reasons outlined at the end of the main text **Methods**. That is, contrary to *current* infections, the probability of having at least one parasite trapped and the total number of parasites trapped in the shell both increase cumulatively with age. Therefore, in the absence of information about age or of a broader set of populations to compare, there is no useful quantitative inference that can be made from these data. Note that encapsulated parasites were nonetheless counted and that these counts are made available along the rest of the raw data (see **Data availability** in main text).

The tables below all exclude the individual that died during the experiment. That snail harboured 102 live nematodes, 0 live trematodes, 0 live mites, had encapsulated 14 nematodes and 0 mites.

**Table S1.** Morph- and population-dependent summary information about nematode prevalence

| shell morph | n | prevalence | 95% interval |
|---|---|---|---|
| **Open habitat** | | | |
| 0 bands | 29 | 0.90 | [0.73; 0.95] |
| 3 bands | 30 | 0.67 | [0.49; 0.80] |
| 5 bands | 30 | 0.40 | [0.26; 0.57] |
| **Shaded habitat** | | | |
| 0 bands | 30 | 0.00 | – |
| 3 bands | 30 | 0.00 | – |
| 5 bands | 30 | 0.00 | – |

**Table S2.** Morph- and population-dependent summary information about nematode intensity of infection

| shell morph | n(infected) | mean intensity | 95% interval | min | max |
|---|---|---|---|---|---|
| **Open habitat** | | | | | |
| 0 bands | 26 | 4.27 | [4.81; 10.63] | 1 | 11 |
| 3 bands | 20 | 5.30 | [3.18; 7.39] | 1 | 12 |
| 5 bands | 12 | 4.33 | [2.27; 6.38] | 1 | 24 |
| **Shaded habitat** | | | | | |
| 0 bands | 0 | – | – | – | – |
| 3 bands | 0 | – | – | – | – |
| 5 bands | 0 | – | – | – | – |

**Table S3.** Morph- and population-dependent summary information about trematode prevalence

| shell morph | n | prevalence | 95% interval |
|---|---|---|---|
| **Open habitat** | | | |
| 0 bands | 29 | 0.14 | [0.07; 0.29] |
| 3 bands | 30 | 0.23 | [0.13; 0.39] |
| 5 bands | 30 | 0.10 | [0.05; 0.23] |
| **Shaded habitat** | | | |
| 0 bands | 30 | 0.13 | [0.05; 0.26] |
| 3 bands | 30 | 0.23 | [0.10; 0.38] |
| 5 bands | 30 | 0.03 | [0.01; 0.14] |

**Table S4.** Morph- and population-dependent summary information about trematode intensity of infection

| shell morph | n(infected) | mean intensity | 95% interval | min | max |
|---|---|---|---|---|---|
| **Open habitat** | | | | | |
| 0 bands | 4 | 1.75 | [1.29; 5.33] | 1 | 4 |
| 3 bands | 7 | 5.71 | [2.52; 9.60] | 1 | 14 |
| 5 bands | 3 | 4.33 | [1.51; 6.97] | 1 | 9 |
| **Shaded habitat** | | | | | |
| 0 bands | 4 | 7.25 | [2.10; 13.04] | 1 | 18 |
| 3 bands | 7 | 7.14 | [2.94; 14.60] | 1 | 28 |
| 5 bands | 1 | 4.00 | [1.13; 14.16] | 4 | 4 |

**Table S5.** Morph- and population-dependent summary information about mite prevalence

| shell morph | n | prevalence |
|---|---|---|
| **Open habitat** | | |
| 0 bands | 29 | 0.00 |
| 3 bands | 30 | 0.00 |
| 5 bands | 30 | 0.00 |
| **Shaded habitat** | | |
| 0 bands | 30 | 0.03 |
| 3 bands | 30 | 0.00 |
| 5 bands | 30 | 0.00 |

**Table S6.** Morph- and population-dependent summary information about mite intensity of infection

| shell morph | n(infected) | mean intensity | min | max |
|---|---|---|---|---|
| **Open habitat** | | | | |
| 0 bands | 0 | – | – | – |
| 3 bands | 0 | – | – | – |
| 5 bands | 0 | – | – | – |
| **Shaded habitat** | | | | |
| 0 bands | 1 | 93.00 | 93 | 93 |
| 3 bands | 0 | – | – | – |
| 5 bands | 0 | – | – | – |

**Table S7.** Morph- and population-dependent summary information about parasite encapsulation in shells

| shell morph | n | proportion with encapsulated nematodes | proportion with encapsulated mites |
|---|---|---|---|
| **Open habitat** | | | |
| 0 bands | 29 | 0.72 | 0.00 |
| 3 bands | 30 | 0.83 | 0.00 |
| 5 bands | 30 | 0.70 | 0.00 |
| **Shaded habitat** | | | |
| 0 bands | 30 | 0.57 | 0.00 |
| 3 bands | 30 | 0.93 | 0.00 |
| 5 bands | 30 | 0.83 | 0.10 |

#### S3 - Pairwise posterior comparisons (multivariate model)

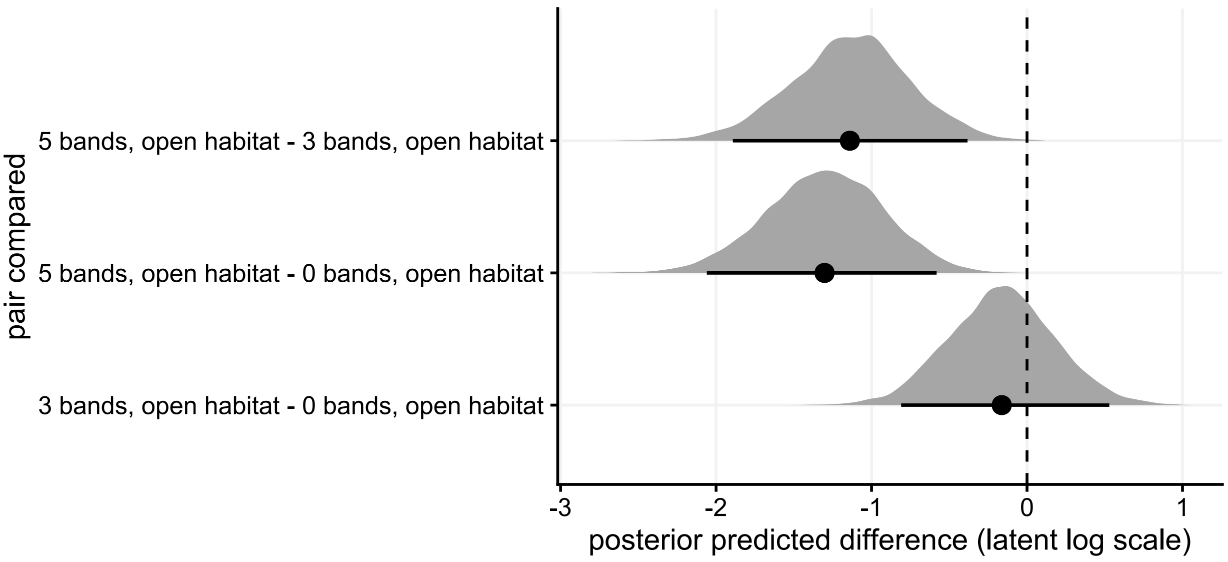


**Figure S1.** Posterior pairwise differences in nematode abundance between snail morphs (on the latent log scale). Reminder: the model for nematode abundance only includes data from the sun-exposed population, since no live nematodes were found in the shaded population.

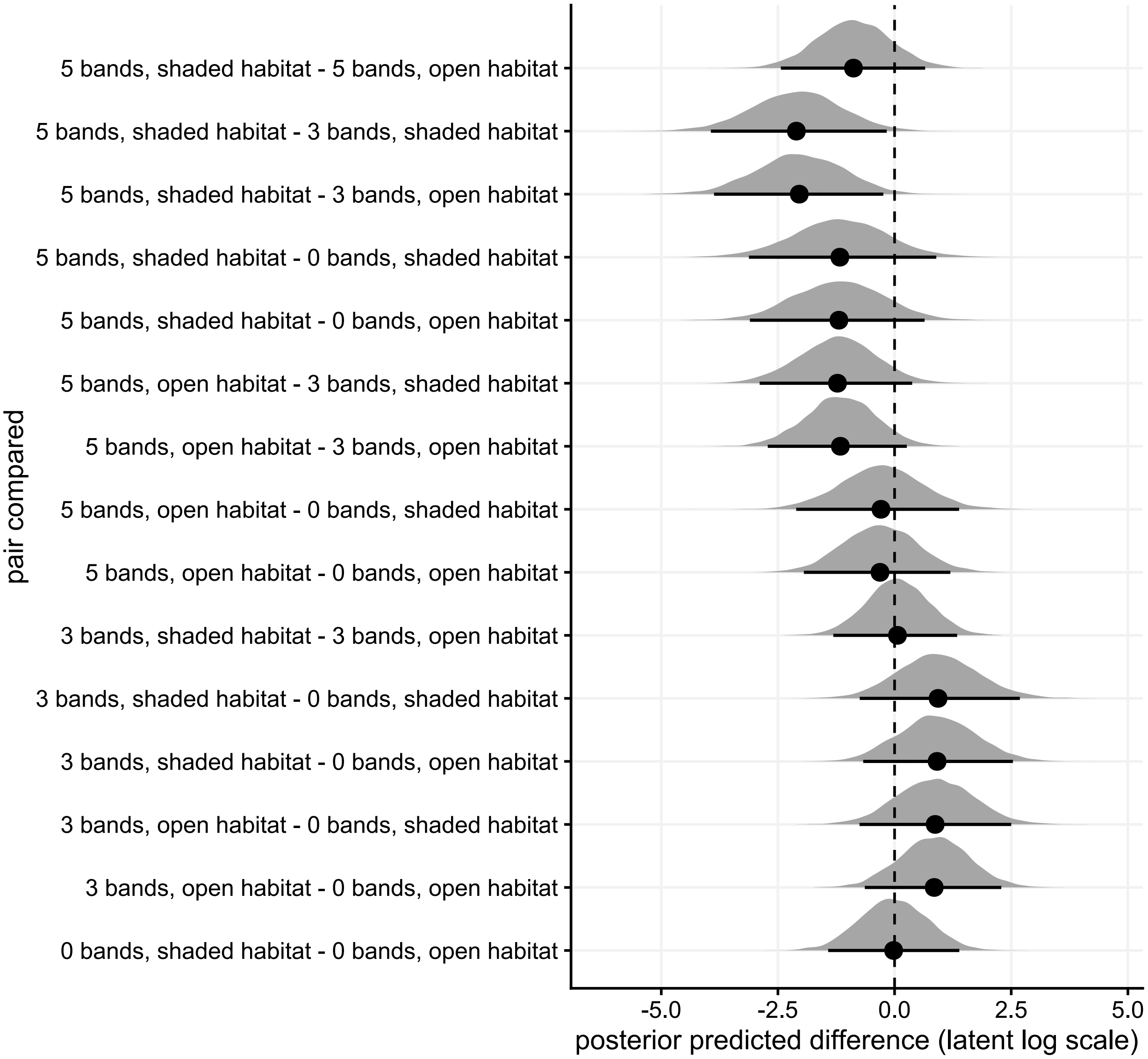


**Figure S2.** Posterior pairwise differences in *Brachylaima* trematode (metacercariae) abundance between snail morph × population combinations (on the latent log scale).

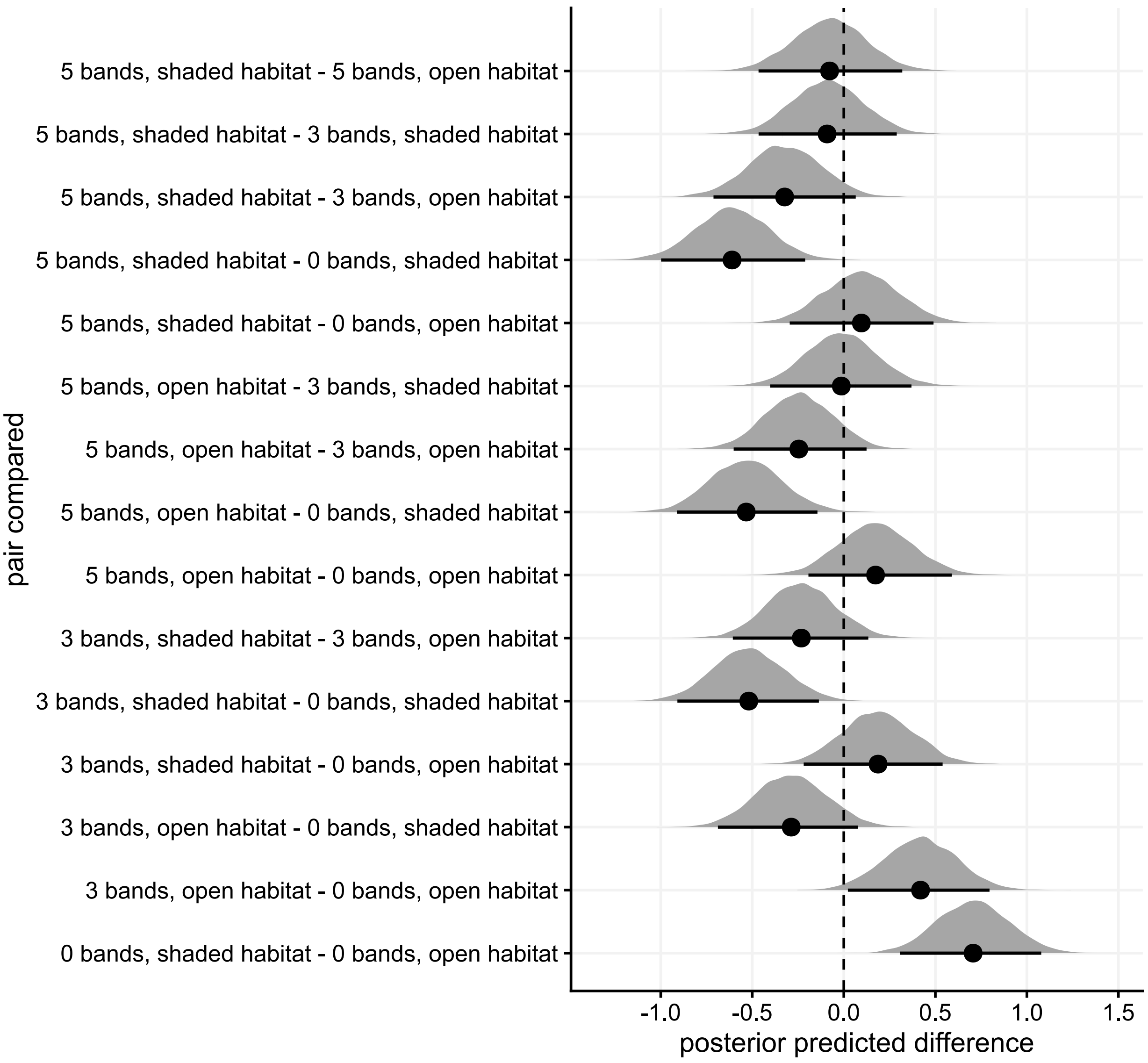


**Figure S3.** Posterior pairwise differences in movement behaviour between snail morph × population combinations. The movement variable is the inverse of the latency to move away (so higher values mean more mobile individuals), and the variable was scaled so differences are in units 1SD.

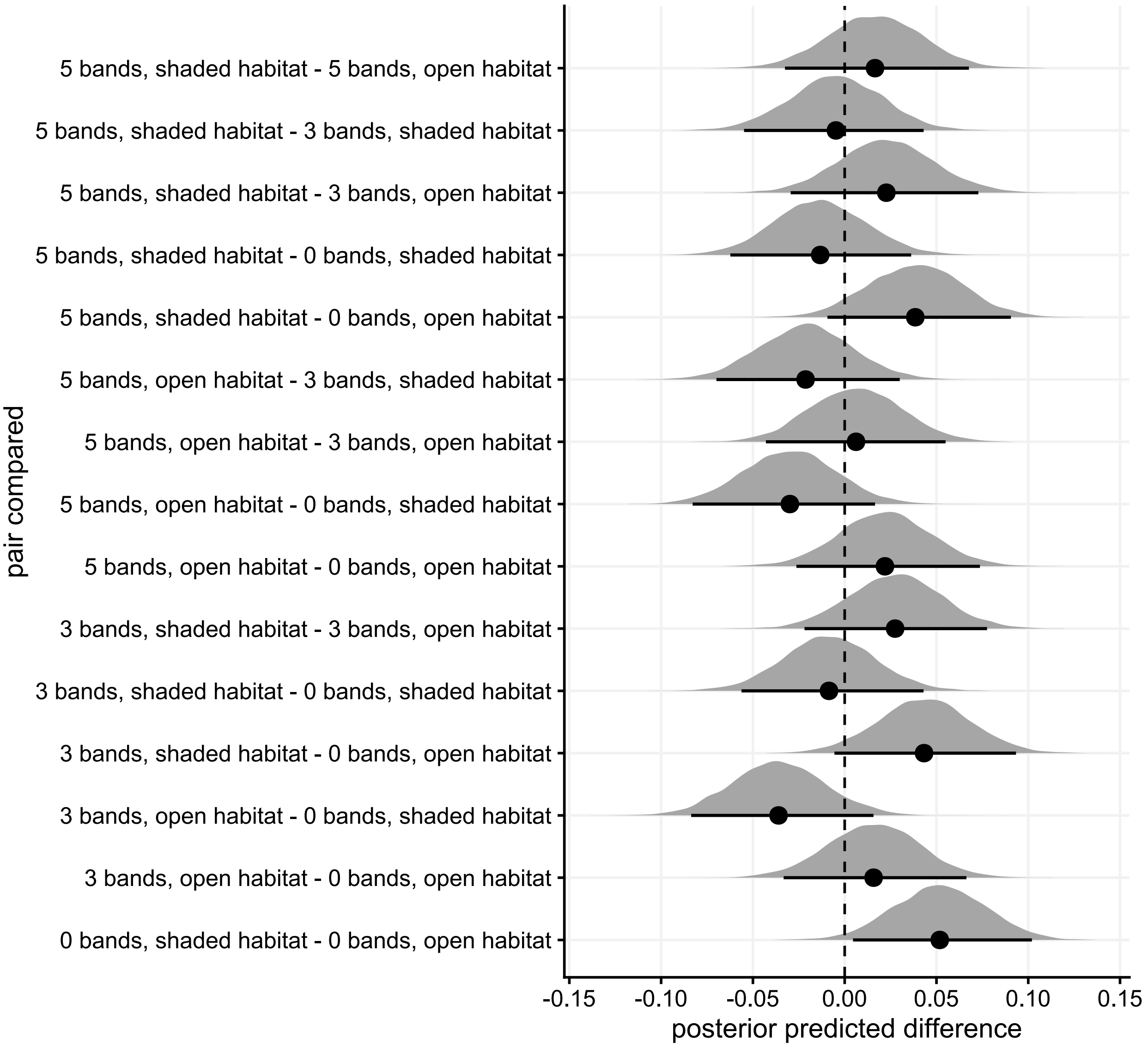


**Figure S4.** Posterior pairwise differences in food intake between snail morph × population combinations.

#### S4 - Posterior proportion of total variance associated with fixed effects vs. random effects in movement and food intake

Both movement behaviour and food intake were observed twice, which allows us to partition variance into among-individual and within-individual components. We show that although in both cases, within-individual/“residual” variation is the dominant component, there is a non-negligible among-individual variance component: both traits are repeatable to some extent (**Supporting Information Figure S5**).

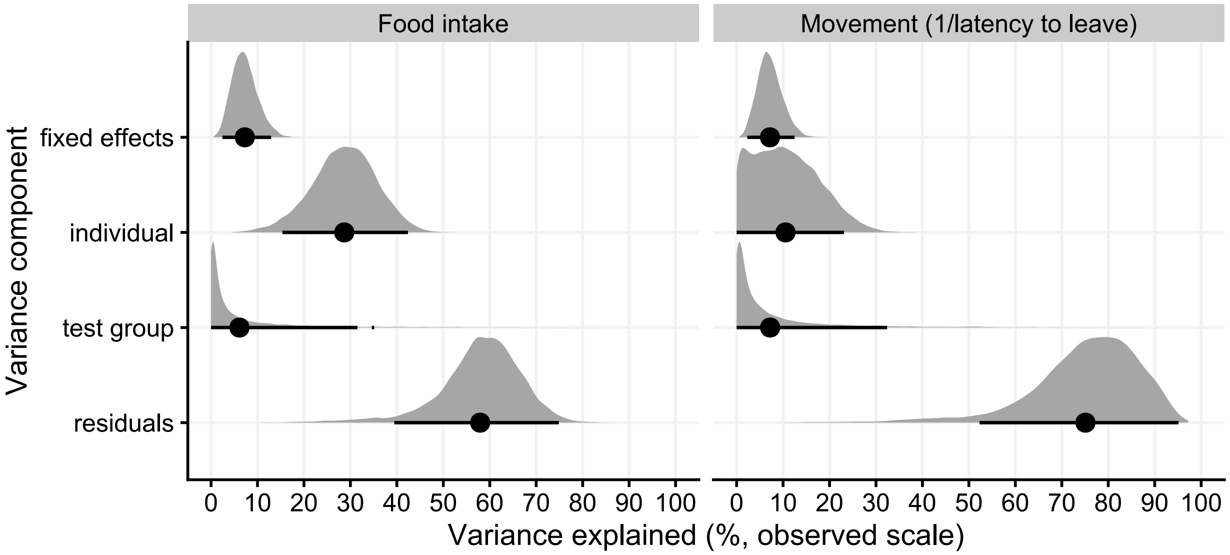


**Figure S5.** Mean (points) and posteriors for the proportion of behavioural variance explained by the different variance components. See **Methods** and **Supporting Information S1** for a description of the model underlying these estimates.

#### References

Bush, A.O., Lafferty, K.D., Lotz, J.M. & Shostak, A.W. (1997). Parasitology meets ecology on its own terms: Margolis et al. revisited. *The Journal of Parasitology* **83**, 575–583.

McElreath, R. (2020). *Statistical rethinking: A Bayesian course with examples in R and Stan.* 2nd edition. Boca Raton, USA: Chapman and Hall/CRC.
